# Circulating Lipids are Associated with PTSD Severity and Predict Symptomatic PTSD in a Cohort of Veterans and Service Members

**DOI:** 10.1101/2025.10.02.680007

**Authors:** Scott Silvey, Ashley Cowart, Jinze Liu, Yang Yue, Jeremy Allegood, Jessica Gill, Christina Devoto, Peter Horvath, Alex Valadka

## Abstract

Post-traumatic stress disorder (PTSD) is characterized by chronic stress, alterations in mood, and avoidance after a traumatic event has occurred. While recovery can occur, many PTSD patients suffer life-long impairments. In combat veterans, high rates of PTSD contribute to increased rates of depression and anxiety and a higher likelihood of suicidal ideation. The lack of biological markers for psychiatric conditions such as PTSD highlights the need for omics-based approaches to diagnosis. Discovery of novel blood-based biomarkers could aid in the development of treatments or therapies, quantify groups for those at the highest risk of adverse events, and provide insight into the molecular underpinnings of PTSD. This study used untargeted lipidomics to analyze 602 circulating lipid species in blood from a cohort of 133 veterans and combat members with varying severities of PTSD. We discovered five circulating lipids, including serum total cholesterol, cholesterol ether (ChE(18:2)), and lipids associated with metabolic dysfunction (cardiolipin : CL(87:7), monolysocardiolipin : MLCL(50:4), and phosphatidylethanolamine plasmalogen/ether lipid : PE(18:1e_20:3)) that correlated significantly with increasing PTSD severity after adjustment for multiple comparisons. Additionally, we showed that symptomatic PTSD patients could be separated from asymptomatic controls using these lipid species. This study contributes to the limited research surrounding the role of circulating lipids in PTSD.

## Introduction

The incidence of post-traumatic stress disorder (PTSD) in United States military personnel exceeds 10 per 1000 and has increased more than tenfold since 2002^1^. PTSD has been estimated to have a prevalence of approximately 15% in deployed U.S. military personnel^2^. A prevalence of 23% has been reported in a meta-analysis of studies of Operation Enduring Freedom/Operation Iraqi Freedom (OEF/OIF) veterans^3^. The importance of PTSD as a widespread mental health problem is not restricted to the military. Approximately 8% of the civilian population will also develop PTSD - and in both military and nonmilitary patients, sequelae of PTSD can include damaged interpersonal relationships, social isolation, divorce, job loss, and suicide.^2^

The diagnosis of PTSD is made clinically according to DSM-5 criteria^4^. However, caveats associated with diagnosis include the fact that symptoms are subjective and self-reported, and they are likely under-reported due to social stigma. Therefore, quantifying severity of PTSD, establishing its chronicity, and tracking response to treatment could be facilitated by identification of objective biomarkers of disease. One potential source of such biomarkers is the serum lipidome.

In recent years, technological developments in mass spectrometry have enabled exploration of thousands of small molecules from a single biological sample. While commonly used clinical tests quantify free fatty acids, triglycerides, and LDL-cholesterol, untargeted lipidomics enables identification of numerous sphingolipids and glycerophospholipids that also circulate in blood while bound to lipoproteins and serum albumin.^5^ Such approaches have revealed the utility of circulating lipids as diagnostic and/or prognostic biomarkers. Most notably, plasma ceramides, specifically the ratio between two distinct ceramide species, have been demonstrated to be more accurate predictors of cardiovascular events such as myocardial infarction and stroke than LDL-cholesterol, a finding that has been exploited for diagnostic use^6^.

While links between plasma lipids and cardiovascular health seem intuitive, use of lipid alterations as indicators of risk for psychiatric disorders is less common. Circulating lipids have been proposed as biomarkers for major depression and bipolar disorder.^7,8^ In PTSD, previous studies have explored metabolomic and lipidomic signatures of disease, finding that levels of compounds linked to metabolic dysfunction and regulation may be altered.^9,10^ However, the literature regarding specific lipid biomarkers in psychiatric disease is scarce, and the overall number of PTSD-related studies is limited. Therefore, we undertook this investigation to determine whether novel circulating lipids could be identified as potentially useful tools for the diagnosis of PTSD and quantification of PTSD severity.

## Methods

The source of the specimens analyzed in this study has been described previously^11^. Briefly, military personnel and veterans were recruited from the community via flyers and advertisements. Data collection took place at two sites, Fort Belvoir Community Hospital and Walter Reed National Military Medical Center. The study was approved by the local institutional review boards. Subjects provided witnessed written informed consent prior to data and sample collection. Participants were active-duty military personnel or veterans, primarily from the OEF/OIF era. Exclusion criteria included severe psychiatric disorders (psychosis, schizophrenia, schizoaffective disorder, bipolar disorder, conversion disorder, or personality disorder). Participants completed a series of questionnaires and provided a blood sample. Samples were collected between 2014 and 2019.

PTSD symptoms were measured using the Posttraumatic Stress Disorder Checklist—Civilian Version (PCLC)^12^. This self-reported measure of PTSD symptoms generates numeric scores ranging from 17 to 85, with scores increasing as severity of PTSD symptoms increases.

Lifetime TBI history was assessed via the Ohio State University Traumatic Brain Injury Identification Method, which is a structured interview administered by a member of the research team^13^. History of TBI (time in years since most recent injury, total number of injuries) was assessed by patient report of injuries to the head that resulted in a period of alteration of consciousness and/or loss of consciousness.

### Untargeted Lipidomics Analysis

Blood samples were stored at -80 degrees C. For processing, samples were thawed on ice. Splash Mix internal standard (Avanti Polar Lipids, Birmingham, AL), was added prior to lipid extraction by the method of Bligh and Dyer^14^. Samples were dried down and redissolved in mobile phase. Drying and reconstituting was accounted for by using internal standards to control for loss during extraction. Untargeted liquid chromatography-tandem mass spectrometry (LC/MS) was performed using a ThermoFisher Q-exactive orbitrap instrument. Data were collected and molecular species were identified by a combination of retention time, molecular mass, and fragmentation pattern using the LipidSearch database (ThermoFisher, USA).

Following pre-processing, we identified 602 lipid species as potential biomarker candidates. Species from both positive and negative modes were identified and combined, and normalized data were used for analysis. Furthermore, lipid intensity values were standardized by centering to the mean and scaling by the standard deviation to increase interpretability.

### Statistical Analysis

Clinical and demographic variables were obtained from the data source, and patient data were summarized. Demographic and clinical variables included patient sex, race, age, body mass index (BMI), combat exposure score, hyperlipidemia status, history of alcohol use, current smoking, TBI history, hypertension, and diabetes status. Demographics and comorbidities were self-reported, and a diagnosis of major depression was made using the PHQ-9 questionnaire. The combat exposure scale (CES) is a self-reported measure ranging from 0-41, that measures various degrees of stress experienced during wartime.

Continuous variables were expressed as the mean□± standard deviation (SD) or median and IQR, and categorical variables were presented as raw values and percentages from the total. We measured the association between total PCLC scores and patient demographic data using Pearson’s correlation test for normally distributed variables, Spearman’s correlation test for non-normally distributed variables, or two-sample *t*-tests for categorical variables. Normality was assessed visually through the use of histograms.

Linear regression tests were conducted, modeling the total numeric PCLC score of each subject at the time of blood collection by each individual lipid intensity value. *P*-values for each resulting model coefficient were obtained, as well as the coefficient estimate itself. Due to previous centering and scaling of lipid intensity values, the β (slope) coefficient for each linear model represented the mean increase in PCLC score for a one-standard deviation increase in lipid intensity. To account for multiple comparisons, raw p-values were adjusted using the method of Benjamini and Hochberg (B/H)^15^, with an overall false discovery-rate (FDR) set to 5%.

After identifying candidate lipids which were statistically significant at B/H-adjusted p<0.05, we fit additional linear regression models for each of these lipids containing all covariates that were significantly associated with PCLC scores. Additionally, major depression showed evidence of a strong association with PCLC scores within our sample. Therefore, we performed a secondary analysis, removing those with a diagnosis of major depression and re-fitting the linear models previously detailed.

### Prediction of Symptomatic PTSD

Next, we aimed to assess the predictive performance of the entire serum lipidome on the classification of symptomatic PTSD versus asymptomatic controls within our cohort. We defined symptomatic PTSD based on total PCLC scores, using previously published definitions (< 25 for asymptomatic, > 50 for symptomatic)^16^. Patients with PCLC scores between 25 and 50 were considered ambiguous cases and thus omitted from this portion of the analysis.

Class separability was visually examined using principal component analysis (PCA)^17^. We then used Orthogonal Projections to Latent Structures-Discriminant Analysis (OPLS-DA) to classify PTSD status^18^. This technique is widely used in the fields of metabolomics and lipidomics. It separates the variation within a sample into predictive and orthogonal (non-predictive) components, with the goal of discriminating between classes while accounting for the inherent biological variability that may occur. Ten-fold cross-validation, stratified by the PTSD status, was used to provide out-of-sample estimates and uncertainties of predictive metrics. We examined balanced prediction accuracy, sensitivity, specificity, positive predictive value (PPV), and negative predictive value (NPV). Individual lipid importance measures were also derived from the model, and the variables of top importance were compared to those obtained in the analysis of continuous PCLC scores.

Statistical analysis was conducted using RStudio version 4.2.3. All hypothesis tests were two-sided. The “ropls” package version 1.30.0 was used to perform OPLS-DA.

## Results

We collected blood samples from *n* = 133 patients. The basic characteristics of our patient sample can be viewed in Table 1. Subjects were majority white (69.2%) and male (78.9%). Around half (45.9%) of patients had a history of alcohol use, and a majority (59.4%) had a diagnosis of major depressive disorder. The median number of TBIs within the cohort was 3 [IQR : 1-5 years], with a median time of 7 years [IQR : 3–11 years] since occurrence of the most recent TBI.

**Table 1:** (*n* = 133) :Cohort Characteristics. ^1^Contained missing values. Only complete cases were considered. ^2^If numeric and normally distributed, obtained using Pearson’s correlation test ; if numeric and non-normally distributed, obtained using Spearman’s correlation test (Total number of TBIs and Time since last TBI) ; if categorical, obtained using two-sample t-test.

| Variable (Units) | Mean (SD), Median (IQR), or N (%) | *p*-value, Association With Total PCLC<sup>2</sup> |
| --- | --- | --- |
| **Demographic/Clinical Characteristics** |  |  |
| PCLC (Total) | 43.23 (16.96) | -- |
| Combat Exposure Score (CES)<sup>1</sup> | 17.84 (10.89) | 0.239 |
| Major Depressive Disorder | 79 (59.4%) | **<0.001** |
| Age | 38.25 (11.36) | 0.271 |
| BMI (kg/m<sup>2</sup>)<sup>1</sup> | 29.12 (4.18) | **0.006** |
| Male Sex | 102 (78.9%) | 0.860 |
| White Race | 92 (69.2%) | 0.097 |
| Total Number of TBIs<sup>1</sup> | 3.0 (1.0 – 5.0) | **<0.001** |
| Time (Years) Since Last TBI<sup>1</sup> | 7.0 (3.0 – 11.0) | **0.007** |
| **Pre-Existing Conditions** |  |  |
| Hyperlipidemia | 25 (18.8%) | 0.409 |
| Hypertension | 32 (24.1%) | 0.114 |
| History of Alcohol Use | 61 (45.9%) | 0.720 |
| Diabetes | 7 (5.3%) | 0.380 |
| Current Smoking | 12 (9.0%) | **0.011** |

The distribution of PCLC scores was approximately normally distributed, with a mean score of 43.23 (SD = 16.96). There was evidence of an association between PCLC scores and body mass index (r = 0.24, p = 0.006), years since last TBI (Spearman’s r = -0.27, p = 0.006), and total number of TBIs (Spearman’s r = 0.46, p < 0.001) (Table 1). Additionally, those with a diagnosis of major depression had significantly higher total PCLC scores (52.34 vs. 29.91, p < 0.001), as did current smokers (54.17 vs. 42.15, p = 0.011). No other covariates showed evidence of association with total PCLC scores.

We identified 602 serum lipids from the untargeted analysis. We tested each individual lipid for a linear association with total PCLC scores among our study sample. The species captured covered a wide range of biological lipids. These included phosphatidylcholines and methyl-phosphatidylcholines (PC/MePC), triglycerides and diglycerides (TG/DG), sphingomyelins (SM), ceramides and glycosphingolipids (Cer/HexCer/Hex2Cer/Hex3Cer), as well as small numbers of acyl-coA carnitines (AcCa), cardiolipins (CL/MLCL), and both lyso-PC and lyso-PE (LPC/LPE) species. Additionally, a moderate number of plasmalogen ether-lipids were identified.

Linear regression tests revealed five lipids with B/H-adjusted p-values < 0.05. The monolysocardiolipin MLCL(50:4) showed the strongest association (β = 7.35), followed by free cholesterol (β = -6.98), both with B/H adjusted p-values < 0.001. A phosphatidylethanolamine plasmalogen/ether lipid : PE(18:1e_20:3) (β = 5.74), cholesterol ester : ChE(18:2) (β = -5.24), and cardiolipin : CL(78:8) (β = 5.15), also had B/H-adjusted p-values under the significance threshold (Table 2). Among other lipids with strong but non-significant associations (raw p-values under 0.01, B/H p-value ≥ 0.05), phosphatidylcholines (PC), Methyl-PC (MePC), diglycerides (DG), and acyl carnitines (AcCa) were generally less abundant as PCLC scores increased, while phosphatidylethanolamines (PE), lyso-PE (LPE), cardiolipins (CL), and SMs were present in higher concentrations as PCLC scores increased. A full summary of those with raw p-values < 0.01 can be found in Table 2. After controlling for BMI, smoking status, years since last TBI, and total number of TBIs, all associations remained statistically significant except ChE(18:2), which was trending (p=0.099, Table 3).

**Table 2:** All lipids with raw *p*-values < 0.01.

| **Lipid Name** | **Regression Coefficient ($\beta$)** | ***P*-Value** | **B/H *P*-Value** |
| --- | --- | --- | --- |
| MLCL.50.4. | 7.35 | <0.001 | **<0.001** |
| Cholesterol (free) | -6.98 | <0.001 | **<0.001** |
| PE.18.1e_20.3. | 5.74 | <0.001 | **0.013** |
| ChE.18.2. | -5.24 | <0.001 | **0.045** |
| CL.78.8. | 5.15 | <0.001 | **0.046** |
| PC.16.0e_18.2. | -4.86 | 0.001 | 0.083 |
| PC.18.1_20.3. | 4.74 | 0.001 | 0.096 |
| TG.12.0_18.2_18.3. | -4.58 | 0.002 | 0.125 |
| PC.16.0e_16.0. | 4.36 | 0.003 | 0.142 |
| PC.18.0e_16.0. | -4.38 | 0.003 | 0.142 |
| AcCa.24.0. | -4.44 | 0.002 | 0.142 |
| PC.16.0_18.1. | -4.43 | 0.002 | 0.142 |
| LPE.18.0. | 4.12 | 0.005 | 0.155 |
| PC.18.1_20.4. | -4.24 | 0.004 | 0.155 |
| MePC.32.1. | -4.19 | 0.004 | 0.155 |
| PC.16.0_16.0. | -4.16 | 0.004 | 0.155 |

| **Lipid Name** | **Regression Coefficient ($\beta$)** | **P-Value** | **B/H P-Value** |
| --- | --- | --- | --- |
| PC.18.0_18.2. | -4.27 | 0.003 | 0.155 |
| PC.38.1. | -4.15 | 0.005 | 0.155 |
| PC.40.8. | -4.13 | 0.005 | 0.155 |
| MePC.34.2. | -4.08 | 0.005 | 0.159 |
| AcCa.16.0. | -4.04 | 0.006 | 0.166 |
| PE.16.0_18.1. | 3.95 | 0.007 | 0.192 |
| PC.16.0_17.0. | -3.89 | 0.008 | 0.194 |
| PE.16.0_20.4. | 3.86 | 0.008 | 0.194 |
| DG.16.0_18.2. | -3.89 | 0.009 | 0.194 |
| DG.34.2e. | -3.89 | 0.008 | 0.194 |
| PC.35.2. | -3.87 | 0.008 | 0.194 |

**Table 3:** Model Table. ^1^Adjusting for BMI, smoking status, number of TBIs, and years since last TBI. ^2^Calculated within those without a diagnosis of major depressive disorder (n = 54, 40.6%).

|  | **Covariate-Adjusted Model<sup>1</sup>** | **Major Depression Excluded<sup>2</sup>** |
| --- | --- | --- |
| **Lipid Name** |  |  |
| MLCL(50:4) | $\beta = 3.33, p=0.017$ | $\beta = 6.33, p<0.001$ |
| Free cholesterol | $\beta = -3.30, p=0.028$ | $\beta = -5.18, p=0.003$ |
| PE(18:1e_20:3) | $\beta = 3.15, p=0.021$ | $\beta = 4.33, p=0.015$ |
| ChE(18:2) | $\beta = -2.34, p=0.099$ | $\beta = -4.31, p=\mathbf{0.016}$ |
| CL(78:8) | $\beta = 2.63, p=\mathbf{0.045}$ | $\beta = 4.58, p=\mathbf{0.009}$ |

Interestingly, all five of the lipids with B/H-adjusted p-values < 0.05 were also significantly associated with a diagnosis of major depression, and the direction of the effect was the same for each lipid. Specifically, free cholesterol and ChE(18:2) were lower in those with a depression diagnosis, while MLCL(50:4), CL(78:8), and PE(18:1e_20:3) were increased (two sample t-tests, all p < 0.001).

In models fit to only the subset of subjects without major depression (n = 54, 40.6%), the isolated effect of each of these lipids on PCLC scores remained consistent and statistically significant. Additional details regarding specific model coefficients can be found in Table 3.

### Prediction of Symptomatic PTSD

We next aimed to assess the predictive performance of the serum lipidome in discriminating between symptomatic post-traumatic stress disorder (total PCLC > 50) and asymptomatic (PCLC < 25) controls. We identified n = 28 (21.1%) asymptomatic controls, and n = 47 with symptomatic PTSD (35.3%) from our cohort. All other patients had PCLC scores which were between 25 and 50, and as a result were not included in this section of the analysis. The PCA plot showed that PTSD-symptomatic and asymptomatic patients displayed a moderate degree of separation. Additionally, non-PTSD control subjects were more tightly clustered, while PTSD-symptomatic subjects displayed more variability. (Fig. 1).

**Figure 1:**
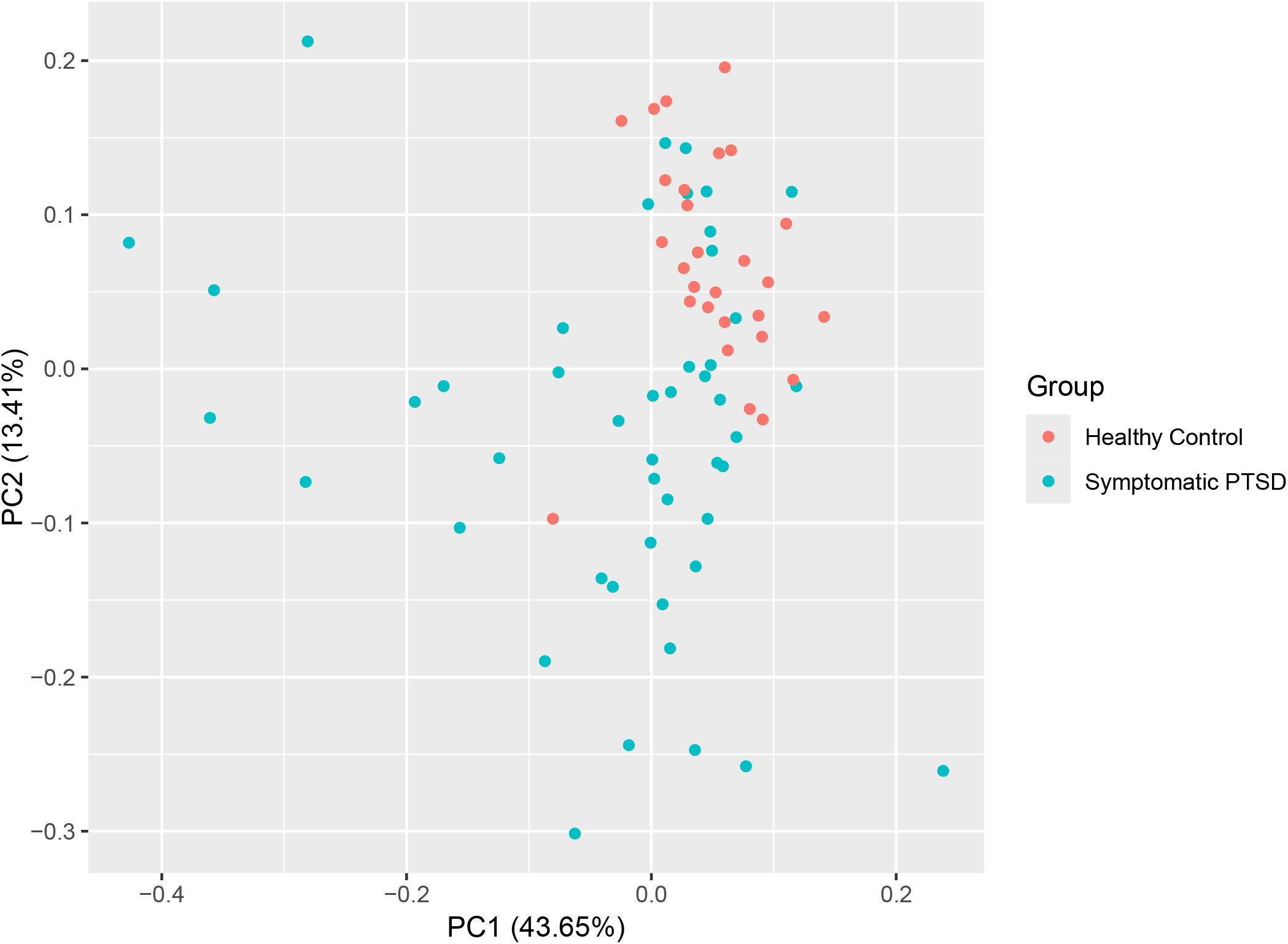
Principal component analysis plot, comparing PTSD-symptomatic versus healthy controls. Points in blue indicate those with PTSD symptoms (PCLC > 50), while points in red indicate healthy controls(PCLC < 25).

Data were split ten folds, maintaining the original proportions of cases versus controls. An Orthogonal Projections to Latent Structures – Discriminant Analysis (OPLS-DA) model was fit to the training set. Cross validation of the model yielded estimates of a balanced accuracy of 68.5% [60.4% - 76.6%], sensitivity of 72.0% [59.3% - 84.7%], and specificity of 65.0% [47.5% - 82.5%], along with positive and negative predictive values of 80.5% [71.5% - 89.5%] and 60.8% [49.2% - 72.5%] respectively. Predictive variable importance measures (VIP) were derived from the full OPLS-DA fit ; the lipids that contributed most to discrimination of PTSD categories were similar to those identified in the univariate analysis, with MLCL(50:4), free cholesterol (ChE), and CL(78:8) having the top VIP measures (Fig. 2). In general, lipids identified to be of top importance in predicting symptomatic PTSD in our cohort were similar to those most associated with PCLC scores in the previous linear regression analysis.

**Figure 2:**
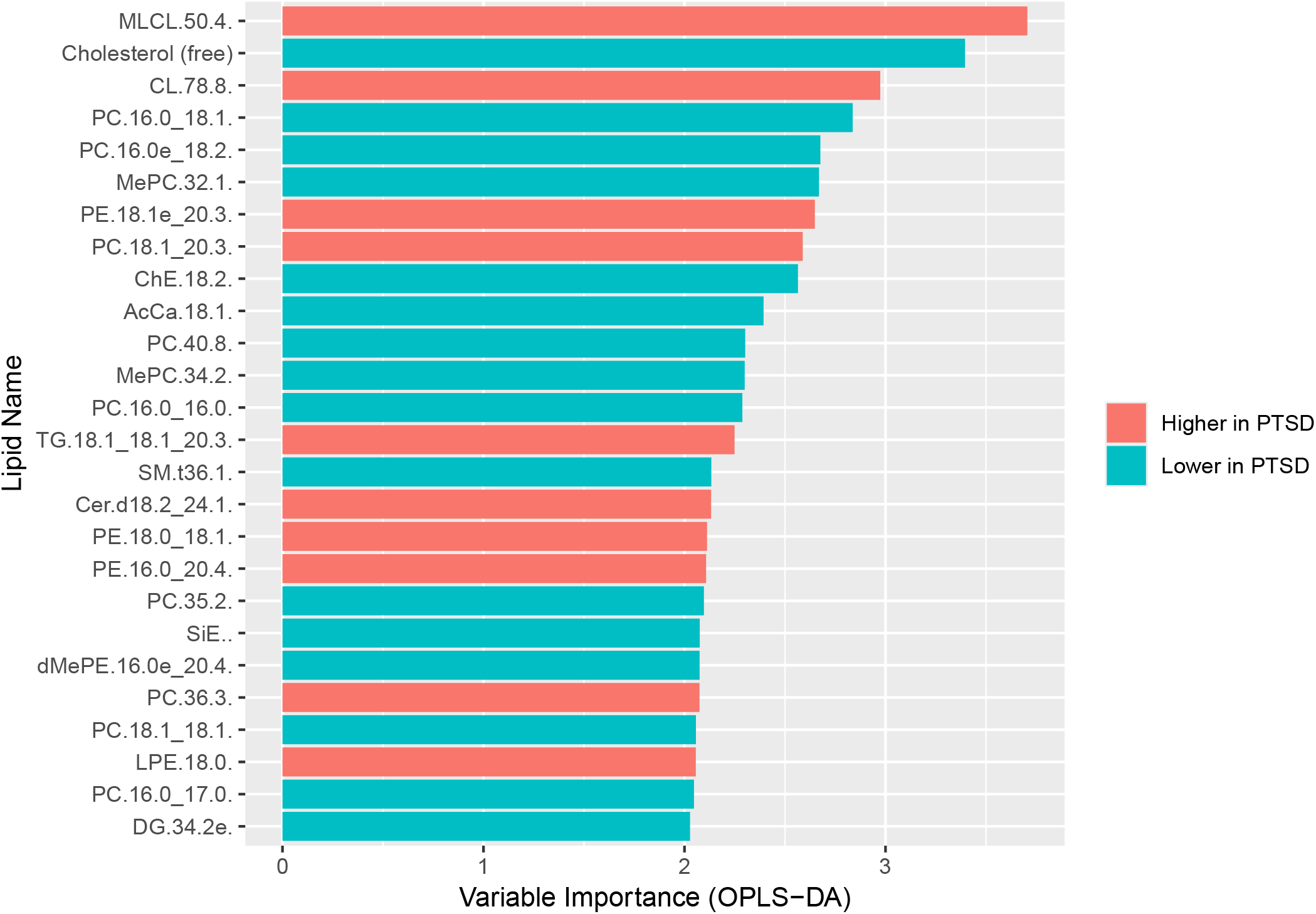
Variable importance plot from OPLS-DA model. Lipids are displayed in decreasing order of importance. The color of the bar indicates how each lipid was altered in PTSD-symptomatic patients (red = higher versus. controls, blue = lower).

## Discussion

A major challenge both with diagnosis and treatment of psychiatric disorders remains the imprecise diagnostic criteria and the heterogeneity of clinical presentation. Through high-throughput, untargeted LC/MS/MS lipidomics analysis of study subject serum, we identified five circulating lipids that were significantly associated with PTSD severity. The lipid with the strongest univariable association and the highest multivariable importance score was MLCL(50:4), which was shown to increase in concentration as PTSD severity increased. Other lipids significantly associated with PCLC scores were cholesterol, ChE(18:2), CL(78:8), and PE(18:1e_20:3). Cholesterol and ChE(18:2) were inversely related to PTSD severity, while CL(78:8) and PE(18:1e_20:3) were positively correlated. After controlling for potential confounders such as TBI-related information, BMI, and smoking status, all lipids remained statistically significant except for ChE(18:2) which was trending. Importantly, despite the high correlation of depression with PTSD, additional stratified analyses showed that the direction and strength of all five lipid effects persisted even in those who did not have an overlapping depression diagnosis. Among other lipids with strong but non-significant associations (raw *p*-values under 0.01, B/H *p*-value ≥ 0.05), phosphatidylcholines (PC), Methyl-PC (MePC), diglycerides (DG), and acyl carnitines (AcCa) were generally less abundant as PCLC scores increased, while phosphatidylethanolamines (PE), lyso-PE (LPE), and SMs were present in higher concentrations as PCLC severity increased.

Next, we created a predictive model that classified symptomatic PTSD in our cohort, which was derived from raw PCLC scores. Although ambiguous patients were excluded, this model was able to correctly identify 72% of symptomatic PTSD cases and 65% of controls correctly when applied to held-out data. Furthermore, if our model predicted a patient to be PTSD-symptomatic, the probability of correct classification was over 80%.

We found evidence of significantly lower serum cholesterol as the symptoms of PTSD severity increased, as well as significantly lower amounts of the cholesterol ester ChE(18:2). Mechanistically, Shrivastava et. al demonstrated that reduction of cholesterol using mevastatin can affect the serotonin system in the brain by reducing ligand-binding and G-protein-coupling abilities of the serotonin receptor. ^19^ Interestingly, the researchers found that actual membrane levels of the serotonin receptor remained unchanged, and that function could be recovered by raising cholesterol levels in the system. Aguiar and Percília showed that treating fish with statins caused increased aggression along with lowered markers of brain serotonin^20^. In humans, numerous studies have found that lower serum cholesterol may be linked to depressive episodes, suicidal or manic states, or other mood disorders^21,22,23^. In PTSD specifically, this association has not been studied extensively, but when it has, there have been mixed results.^24^ There is also strong evidence of an overlap between PTSD and other psychological disorders like major depression^25^. Additionally, comorbidities such as heart disease, obesity, and diabetes - which are themselves associated with *high* serum cholesterol - are also linked to PTSD and may be attributed to both lifestyle and genetic factors^26,27^. In general, the interplay between PTSD, physical comorbidities, and serum cholesterol requires more research, but if validated may be a promising measure to quantitatively identify those most at risk for more severe outcomes such as suicidal thoughts and behaviors or violence.

Among the other identified lipids, MLCL(50:4) and CL(78:8) were shown to be increased in those with higher PTSD severity. These are cardiolipin species, which are strongly linked to mitochondrial function^28,29,30^. Although cardiolipin is mainly found in the mitochondria of eukaryotic cells, it has been shown that these species are also abundant in the blood and are enriched in low-density lipoprotein (LDL) .^31^ In human disease, MLCL and other species have been shown to acutely rise after traumatic brain injury due to cardiolipin oxidation, which may be causative of brain damage.^32,33^ In our study, the effects of MLCL(50:4) and CL(78:8) on PTSD severity were consistent, regardless of the time since each patient’s last TBI or the total number of TBIs per-patient. A targeted lipidomic approach is necessary to validate and study specific mechanisms, but our results suggest that a chronic increase in these species, potentially caused by oxidative stress in the brain, may be a biological indicator of more severe PTSD symptoms.

Finally, we observed an increase in PE(18:1e_20:3), which is a plasmalogen ether-lipid. These types of lipids have also been linked to metabolic dysfunction and cognitive decline, although the biological mechanism behind ether-lipids and disease is unclear.^34,35^

Limitations of this study include its small sample, cross-sectional, and retrospective nature, which may not include or highlight confounders of the relationship between these lipids and PTSD severity. Additionally, there could be a potential lack of generalization to a non-veteran population and a single measure of PCLC alone as a self-reported measure of symptom severity may be more variable than repeated measurements of this test in conjunction with other questionnaires. By using untargeted lipidomics, we were able to examine a much larger number of lipid species than a targeted approach, but the confidence behind the exact chemical structures of some identified lipids is less reliable. Finally, we did not have access to specific medication data such as statins, which affect lipid levels, although these were unlikely to affect our results as PCLC scores were not associated with the rates of hyperlipidemia, diabetes, or hypertension in this cohort. Potential avenues for future research include targeted lipidomics to validate the biomarkers that we observed, risk stratification based on biomarker levels, as well as comparison to a non-veteran control group. We hope that this study, as well as future research, will contribute to the limited literature surrounding novel diagnostic and therapeutic approaches within the context of precision psychiatry and stress disorders.

## Acknowledgements

Supported by a generous gift from the David P. Reynolds DRB Fund

## Data Availability

Can be shared if requested post-acceptance.

